# Decrypting protein surfaces by combining evolution, geometry and molecular docking

**DOI:** 10.1101/507095

**Authors:** Chloé Dequeker, Elodie Laine, Alessandra Carbone

**Affiliations:** Sorbonne Université, CNRS, IBPS, Laboratoire de Biologie Computationnelle et Quantitative (LCQB), 75005 Paris, France; Institut Universitaire de France (IUF), Paris, France

## Abstract

The growing body of experimental and computational data describing how proteins interact with each other has emphasized the multiplicity of protein interactions and the complexity underlying protein surface usage and deformability. In this work, we propose new concepts and methods toward deciphering such complexity. We introduce the notion of interacting region to account for the multiple usage of a protein’s surface residues by several partners and for the variability of protein interfaces coming from molecular flexibility. We predict interacting patches by crossing evolutionary, physico-chemical and geometrical properties of the protein surface with information coming from complete cross-docking (CC-D) simulations. We show that our predictions match well interacting regions and that the different sources of information are complementary. We further propose an indicator of whether a protein has a few or many partners. Our prediction strategies are implemented in the dynJET^2^ algorithm and assessed on a new dataset of 262 protein on which we performed CC-D. The code and the data are available at: http://www.lcqb.upmc.fr/dynJET2/.

**Author summary:** The multiplicity and versatility of protein interactions make protein surfaces complex biological objects. For instance, a protein may interact with several partners at different moments via the same region at its surface, or it may use several regions, with different shapes and evolutionary origins, to establish different types of interactions. In addition, interfaces can re-adjust depending on environmental conditions. In this work, we introduce the notion of ‘‘interaction region”, in contrast to ‘‘interaction site”, which accounts for the interface variability coming from molecular flexibility and for the multiple usage of the protein surface by several partners. Moreover, we use four biologically meaningful descriptors to delineate protein surface patches. Three of them, namely evolutionary conservation, physico-chemical composition and local geometry, are computed for a given protein, without any knowledge on its potential partners. The fourth property is inferred from the behavior of the protein with respect to other proteins, partners or not, in docking simulations. We show that predicted patches match well with known interacting regions, that the four descriptors are complementary and that they enable capturing most of the signal relevant to protein interactions. We further exploit the way patches are grown to precisely localize interaction regions, and to estimate whether a protein have a few or many partners.

## Introduction

Proteins are main actors in biological processes and a detailed description of their interactions is expected to provide direct information on these processes and on the way to interfere with them [1]. Our knowledge of protein-protein interaction (PPI) networks [2] is largely incomplete, since the experimental assessment of all possible interactions of a protein is very challenging [3, 4]. To overcome this limitation, recent efforts have been invested in the integration of direct and indirect experimental evidence and of computational predictions to better describe PPIs at the genome scale [5, 6, 7, 8, 9, 10, 11]. These efforts have revealed the complexity and multiplicity of PPIs. A protein may interact with several partners at the same time - each partner binding to a different site at its surface, or its surface may present a shared binding region that will be used by different partners at different moments of its lifetime. It is estimated that as much as 75% of the surface could potentially be used for PPIs [12]. In this context, there is a need for the development of tools able to decrypt protein surfaces at the residue level. A comprehensive description of protein surfaces and a precise identification of the residues involved in interactions are mandatory to identify cellular partners at large scale [11] and design drugs modulating PPIs [13]. Moreover, characterizing protein surfaces’ properties may inform us on the number of partners a protein may have, and thus on the role of that protein in the cell.

Evolutionary, physico-chemical and geometrical properties have been shown to be relevant to PPIs [14, 15, 16, 17, 18, 19, 20, 21, 22, 23, 24, 25], and, based on them, in the past 20 years, a number of tools have been developed to predict interacting surfaces [26, 27, 28, 29, 30, 31, 32, 24, 33] (see [25, 34] for surveys). Although some of these tools achieve very high accuracy against subsets of known experimental binding sites, their predictions are generally much smaller than the expected interacting surface size [12]. Moreover, many tools do not propose sites but rather evaluate the probability of a residue to be involved in interactions. An orthogonal approach consists in exploiting molecular docking calculations. Docking methods were originally designed to predict the structure of a complex starting from the known structures of its components. Candidate conformations are evaluated based on properties reflecting the strength of the association, *e.g*. shape complementarity, electrostatics, desolvation, conformational entropy. By deriving statistics from the generated conformational ensemble, one can estimate the propensity of each protein surface residue to be found at a docked interface and use these propensities to identify binding sites [35]. This has been realized in single docking studies [36, 37, 38, 39, 40], where two proteins known to interact are docked to each other, in arbitrary docking studies [41], where proteins from a benchmark set are docked to arbitrarily chosen proteins, and in complete cross-docking studies [42, 11, 43, 44, 45], where all versus all docking is realized on a given dataset.

In the present study, we combine these different types of information to decipher the complexity of protein surfaces and give clues about the many interactions a protein may have (Fig. 1). Given a protein, we predict patches at its surface based on some intrinsic properties of that surface and on properties inferred from the behavior of the protein with respect to others in docking calculations. We assess our predictions against a new set of experimentally known functional interfaces, detected at the surface of 262 proteins and of their close homologs. We demonstrate that considering only one single complex for a given protein leads to underestimate the proportion of its surface involved in functional interactions and to the incorrect assessment of protein interface prediction algorithms. To cope with this issue, we introduce the new concept of interacting region (IR) as a protein surface region used by one or several partners. IRs are defined by merging overlapping interacting sites (IS) extracted from different protein complex structures. We show that our predictions better match IRs compared to ISs and capture the interface variability induced by molecular flexibility. Our approach includes sequence-based analysis, which allows the detection of signals even when the interface is ‘‘hidden”. Interestingly, we highlight a few cases where docking enables unveiling interfaces that could not be detected otherwise. We further exploit the way in which our predicted patches are grown, starting from a seed that is progressively extended. Specifically, we demonstrate that predicted patches’ seeds can be used to localize IRs with high precision and to determine whether a protein has a few or many partners.

**Figure 1:**
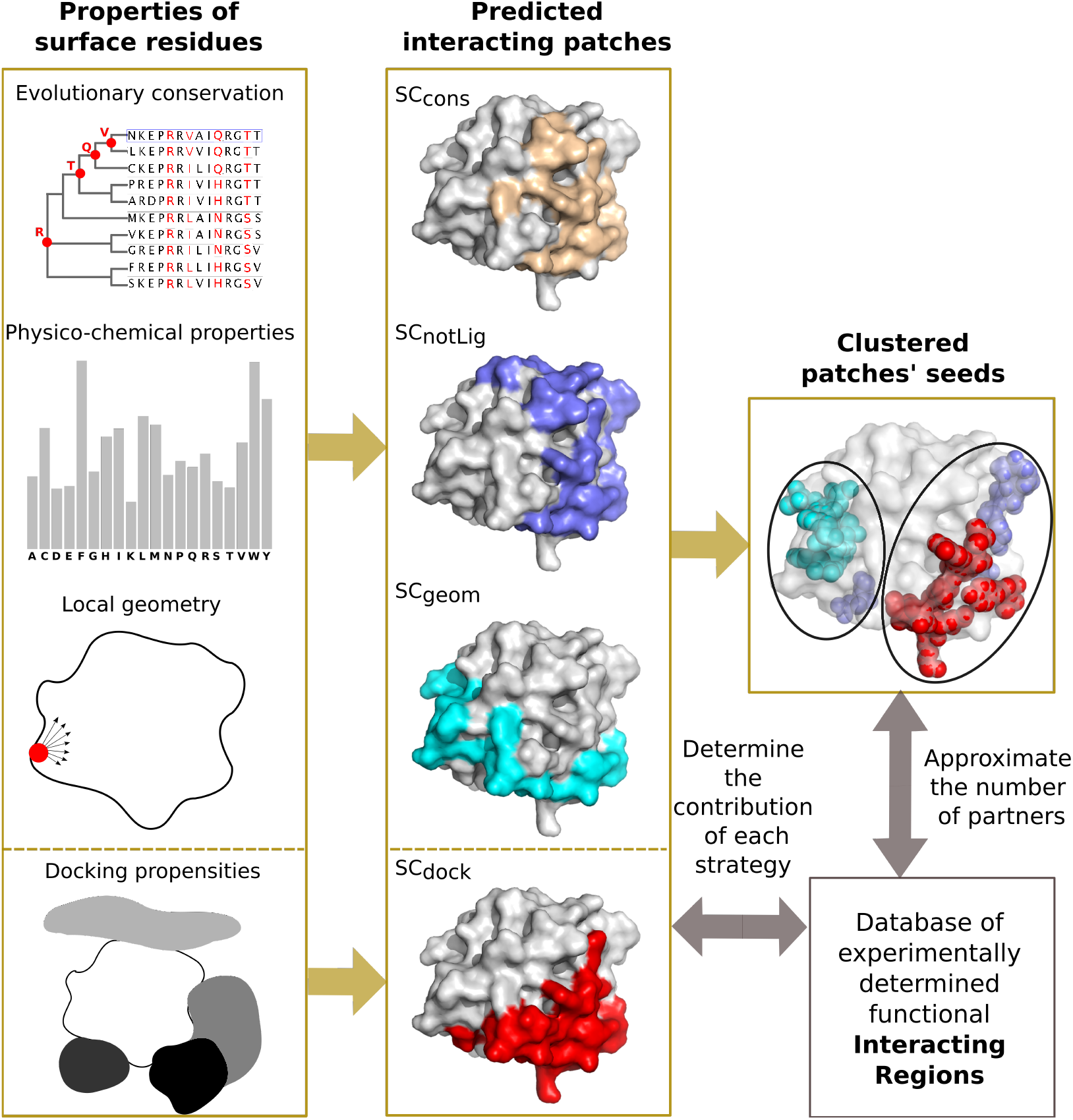
Schematic representation of our workflow. We consider four residue-based properties (left panel), namely evolutionary conservation, amino acid propensities to be found at an interface, local geometry and propensities to be found in docked interfaces. We predict interacting patches at the surface of proteins by using four different strategies: SC_*cons*_, SC_*notLig*_ and SC_*geom*_ combines the first three properties, while SC_*dock*_ relies exclusively on the fourth property. We compare the predicted patches with a set of experimentally determined functional interacting regions. We analyze and cluster the predicted patches’ seeds, from which they were grown, to precisely localize interacting regions and infer the number of partners used by each region.

We provide sets of experimentally known interaction sites and regions and complete cross-docking results for our dataset of 262 proteins, along with a computational tool, called dynJET^2^, for predicting interacting patches based either on protein sequence and structure analysis or on any pre-computed residue based property. All data and implemented code are available at: http://www.lcqb.upmc.fr/dynJET2/.

## Results

### From interacting sites to interacting regions

Our analyses were performed on a set of protein complex crystallographic structures, which we call `P-262`, involving 262 protein chains (see Materials and Methods and S1 Table). From these experimental structures, we defined two sets of functional interfaces, `PPI-262` and PPI-262_ext_ (see Materials and Methods and S1 Fig). `PPI-262` comprises 329 ISs, where each IS corresponds to one functional interaction described by one structure. This classical definition of a protein interacting site is very restrictive and does not account for the interface variability that may come from structure ensembles. Indeed, the definition of the interface between two given proteins may vary from one structure to another, depending on the crystallization conditions, on the quality of the data/model and/or on the inherent flexibility of the assembly. What is more, the notion of IS masks the complexity of protein surface usage by multiple partners. This motivated us to define the new concept of IR, obtained by merging overlapping ISs (≥ 60% overlap). Based on the observation that functional interfaces are conserved across closely related homologs [46], we collected all functional ISs involving the query proteins from `P-262` or their close homologs (≥ 90% sequence identity) from the Protein Data Bank [47] (PDB). This amounted to 23 642 ISs, which were merged into 370 IRs to define our second “extended” dataset, PPI-262_ext_. The two examples in Fig. 2 illustrate the complexity of the experimental interaction surfaces in our datasets. Binding sites may be disjoint, overlapping or included in others (Fig. 2, on the left), and they may be defined by the interaction with several proteins or peptides (Fig. 2, on the right). The two examples show 5 ISs (3 on the left and 2 on the right), which were merged into 3 distinguished IRs (2 on the left and 1 on the right, contoured by thick forest green lines). In all those cases, the IRs result from the merging of ISs that represent binary interactions with different partners. In addition, IRs may also be defined from several ISs representing a single interaction, but whose binding mode slightly differs from one PDB structure to another (see below).

**Figure 2:**
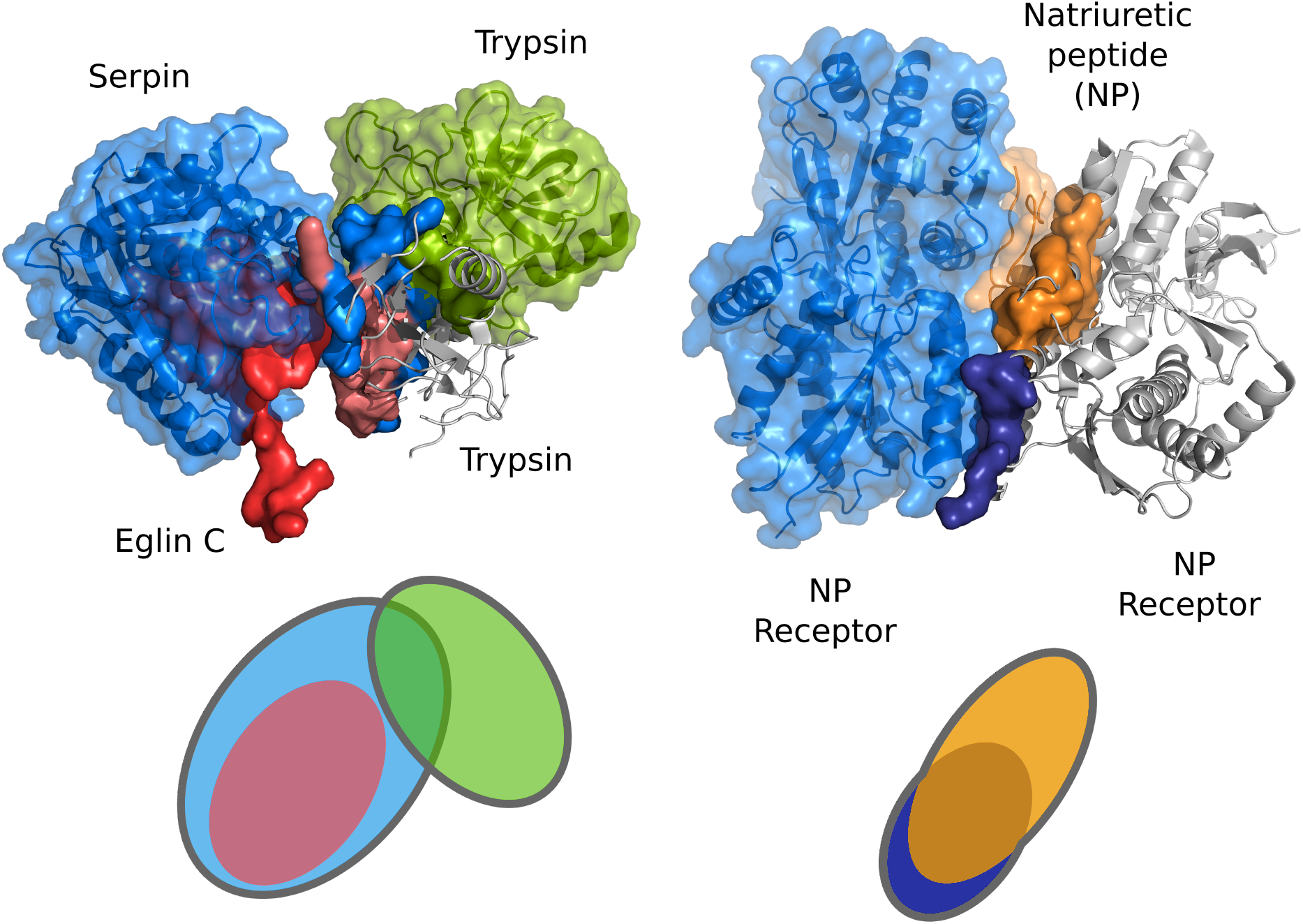
Two examples of the usage of the protein surface by different partners. The query proteins are displayed as grey cartoons, their interacting sites as opaque colored surfaces and their partners as colored cartoons and transparent surfaces. Left: trypsin (1ezx_C, in grey) interacts with itself (5gxp_B, in green), with serpin (1ezx_A, in blue) and with eglin C (4b2b_B, in red). The 3 corresponding ISs lead to the definition of 2 IRs, as depicted on the schema at the bottom, where each IR is contoured by a thick grey line. Right: the natriuretic peptide receptor forms a homodimer (1yk1_A, in grey, and 1yk1_B, in blue) to bind its substrate (1yk1_E, in orange). The 2 ISs detected at the surface of one receptor monomer (1yk1_A, in grey) are merged into an IR.

### Prediction of interacting patches

We predicted interacting patches at the surface of the proteins from `P-262` by relying on four residue properties (see Materials and Methods for precise definitions): evolutionary sequence conservation inferred from the analysis of homologous sequences, physico-chemical properties expected at the interface based on experimentally known complex structures, local geometry computed on the protein 3D structure, and propensities to be found at docked interfaces inferred from complete cross-docking calculations (Fig. 1). The first three properties are used to derive three different scoring strategies (SC_*cons*_, SC_*notLig*_ and SC_*geom*_) aimed at identifying different types of protein-protein interfaces (see Materials and Methods and [24]). Each SC explicitly describes the role of each one of the properties with respect to the expected support-core-rim structure of interacting sites [48]. Evolutionary sequence conservation is used (SC_*cons*_) to target very conserved residues on the protein surface and detect most types of interfaces. Local geometry is combined with evolutionary conservation (SC_*notLig*_) to specifically discriminate protein-protein interfaces from small-ligand binding pockets, the latter being usually more deeply buried. SC_*cons*_ and SC_*notLig*_ are both designed to target conserved sites, and as a consequence their predictions often overlap substantially [24]. By contrast, physico-chemical properties are combined with local geometry (SC_*geom*_) to capture highly protruding interfaces not conserved through evolution. The fourth docking-based property is used exclusively in a fourth strategy, SC_*dock*_ (see Materials and Methods). This property reflects the propensity of each protein residue to bind partners and non-partners in docking calculations. To evaluate docking conformations, we used a coarse-grained empirical energy function comprising a Lennard-Jones potential for van der Waals interactions and a Coulomb potential for electrostatics [49]. The four SCs are implemented in dynJET^2^, an upgraded version of the JET^2^ method [24].

### Estimation of the protein surface involved in functional interactions

We used both experimental interfaces and predicted patches to estimate the proportion of protein surface involved in functional interactions. On average, experimental ISs from `PPI-262` cover 29% of the protein surface (Fig. 3a). Hence, by looking at this dataset, one may infer that the residues involved in functional interactions generally represent less than a third of the protein surface. However, when looking at PPI-262_ext_ (Fig. 3b), which comprises experimental IRs defined from close homologs, the coverage increases up to 48%. Moreover, a significant number of proteins (32) have their surface completely or almost completely covered by functional interactions (coverage ≥80%). This suggests that most of the proteins from P-262 engage in multiple interactions with different partners.

**Figure 3:**
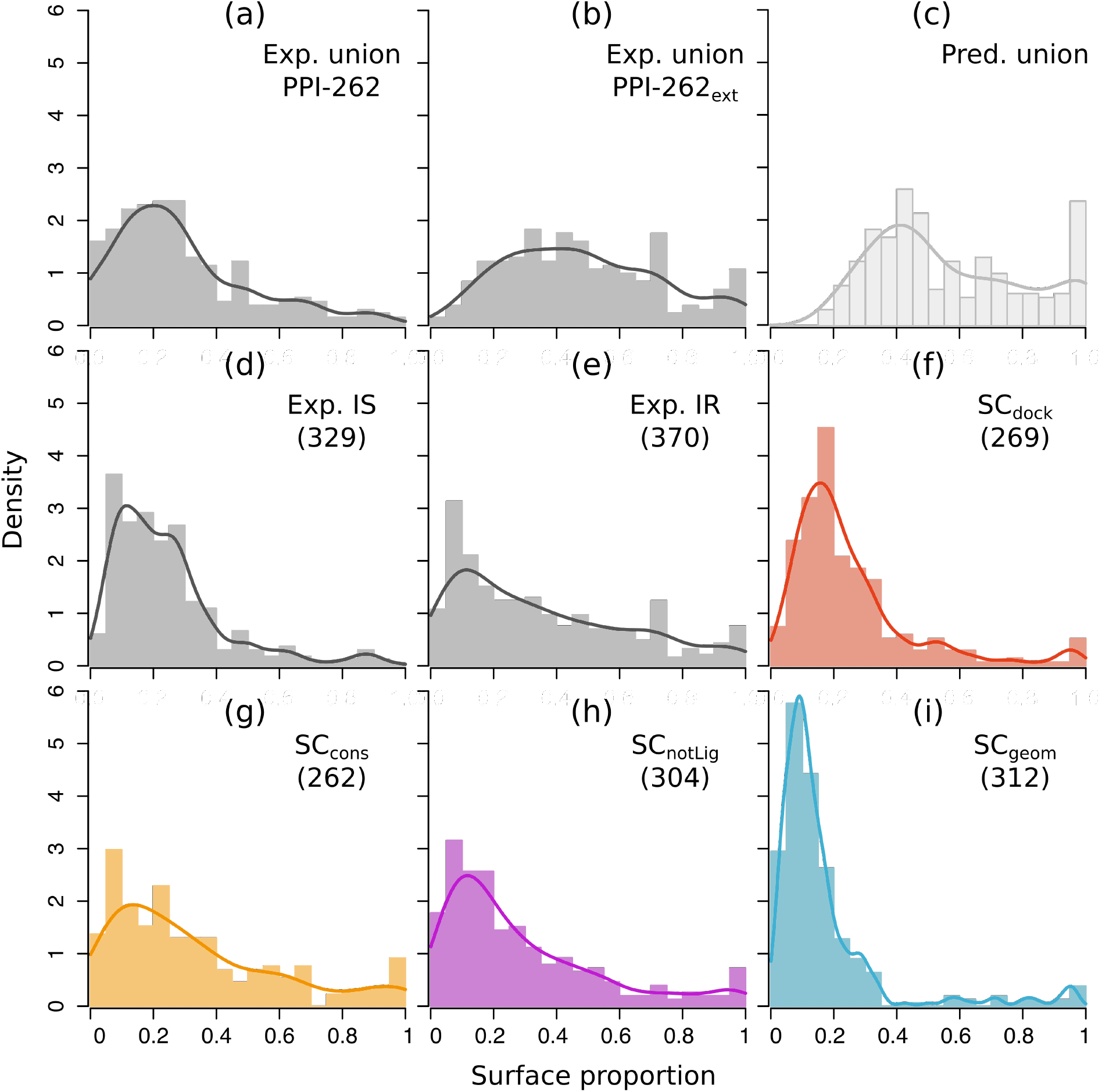
Proportion of protein surface covered by experimental interfaces and predicted patches. Distribution are reported for: (**a**) the union of ISs from `PPI-262`, (**b**) the union of IRs from PPI-262_ext_, (**c**) the union of patches predicted by dynJET^2^, (**d**) individual ISs from `PPI-262`, (**e**) individual IRs from PPI-262_ext_,(**f-i**) individual patches predicted by each SC. The union of ISs, IRs or predicted patches is realized for each protein. Notice that the sizes of the predicted patches do not add up when considering their union, since several of them overlap.

The estimation provided by the union of predicted patches is slightly higher, 56% on average (Fig. 3c). The associated distribution resembles that of the union of experimental IRs from PPI-262_ext_ (compare Fig. 3c with 3b), except for two notable differences at the extremities. This observation is statistically supported by a Mann-Whitney U-test: while the full distributions are different (p-value=0.002), the truncated distributions (without values below 20% and above 95%) are indistinguishable (p-value=0.37). The differences at the extremities are the following: the minimum coverage is higher for predictions than for experimental interfaces (18% versus 6.2%), and there are more proteins completely or almost completely covered (≥ 80%) by predictions than by experimental interfaces. The first difference can be explained by the specifics of dynJET^2^ clustering algorithm, which discards very small predictions (see Materials and Methods and [23]). The second difference suggests that all functional interfaces have not been yet experimentally characterized.

When looking at individual patches instead of their union, we found that patches predicted from docking (SC_*dock*_) display sizes similar to those of experimental ISs (compare Fig. 3d and 3f). Both types of interfaces represent about one quarter of the protein surface, on average. By contrast, conserved (SC_*cons*_, SC_*notLig*_) predicted patches are bigger, covering about one third of the protein surface, on average (Fig. 3g,h). Their size distributions are similar to that of experimental IRs (compare with Fig. 3e). These three types of interfaces are highly variable, with standard deviations in the [24 – 28]% range. Finally, patches predicted based on local geometry (SC_*geom*_) are the smallest (Fig. 3i), representing 16% of the protein surface.

### Assessment of the predictions and contribution of each SC

The identification of a protein’s set of interacting residues is important to understand the determinants of molecular association. For each protein, we compared the union of all predicted patches with the union of all ISs (respectively IRs) from `PPI-262` (resp. PPI-262_ext_). To do so, we relied on the F1-score, which reflects the balance between precision (or positive predictive value) and recall (or sensitivity). The average F1-score on `PPI-262` is 0.41 ± 0.24 and it increases up to 0.57 ± 0.19 on PPI-262_ext_ (Fig. 4a). This increase reflects a global shift of the F1-score distribution toward higher values (`p-value=`10^-4^ with the Mann-Whitney U test). The proportion of proteins with very good predictions (F1-score > 0.6) increases from 18% to 46%, while that of proteins with very poor predictions (F1-score < 0.2) drastically reduces from about one quarter to 4%. These results highlight the importance of considering all available experimental information to properly evaluate protein interface predictions. Predicted residues that would be considered as false positives when looking at the restricted dataset, `PPI-262`, are actually involved in interactions with other partners, as revealed by the extended dataset, PPI-262_ext_.

**Figure 4:**
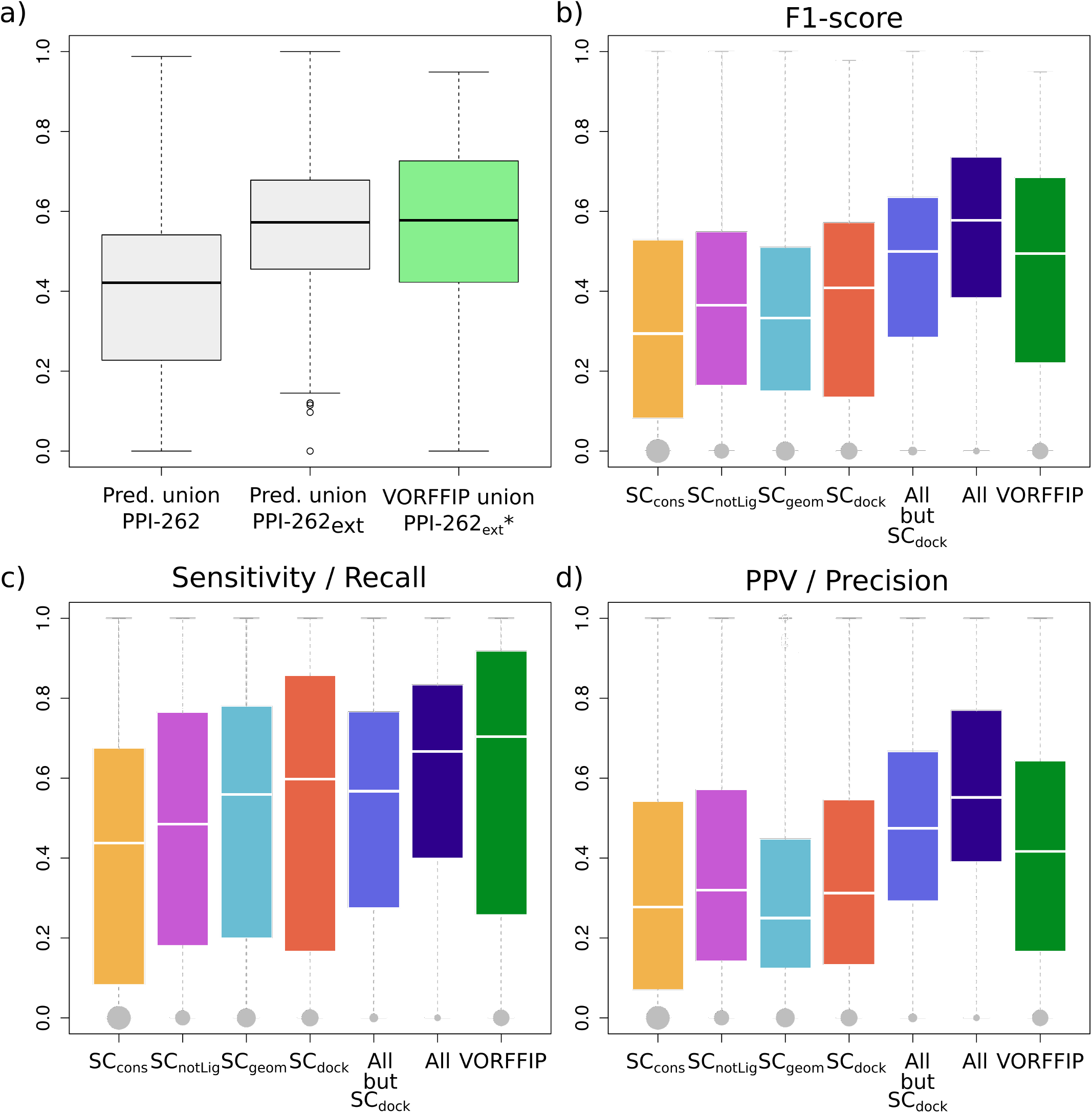
Agreement between experimental interfaces and predicted patches. (a) Distribution of F1-scores computed for the union of dynJET^2^ predictions against the union of ISs from `PPI-262` (in light blue) and of IRs from PPI-262_ext_ (in dark blue), and for the union of Multi-VORFFIP predictions against PPI-262_ext_*, a subset from PPI-262_ext_ involving 252 protein chains (in green). (b-d) Agreement between predicted patches and experimental IRs from PPI-262_ext_. For each IR, the best-matching patch or combination of patches predicted by the SC(s) indicated in x-axis is retained. The performance measures are the following: (b) F1-score, (c) sensitivity (recall), (d) positive predicted value (precision). The sizes of the grey dots are proportional to the number of IR that could not be detected at all.

We further investigated to what extent the partitioning of protein surfaces into patches predicted by the different SCs matches experimental IRs (Fig. 4bcd). None of the SC is sufficient on its own to detect all IRs (Fig. 4c, in orange, purple, cyan and red). This observation is also illustrated by the two examples of Figs 5a and 5c, where several SCs are necessary to capture the entirety of the experimental signal. Combining SC_*cons*_, SC_*notLig*_ and SC_*geom*_. enables increasing the average F1-score by about 0.1 compared to individual SCs, and drastically reducing the number of completely missed IRs to only 28 over 370 (7.6%, Fig. 4b, in marine). This is indicative of the complementarity of the three SCs in their coverage of the protein surface, as already observed in [24]. Accounting for SC_*dock*_ patches further enhances the quality of the predictions up to an average F1-score of 0.54 (compare boxplots in marine and darkblue).

**Figure 5:**
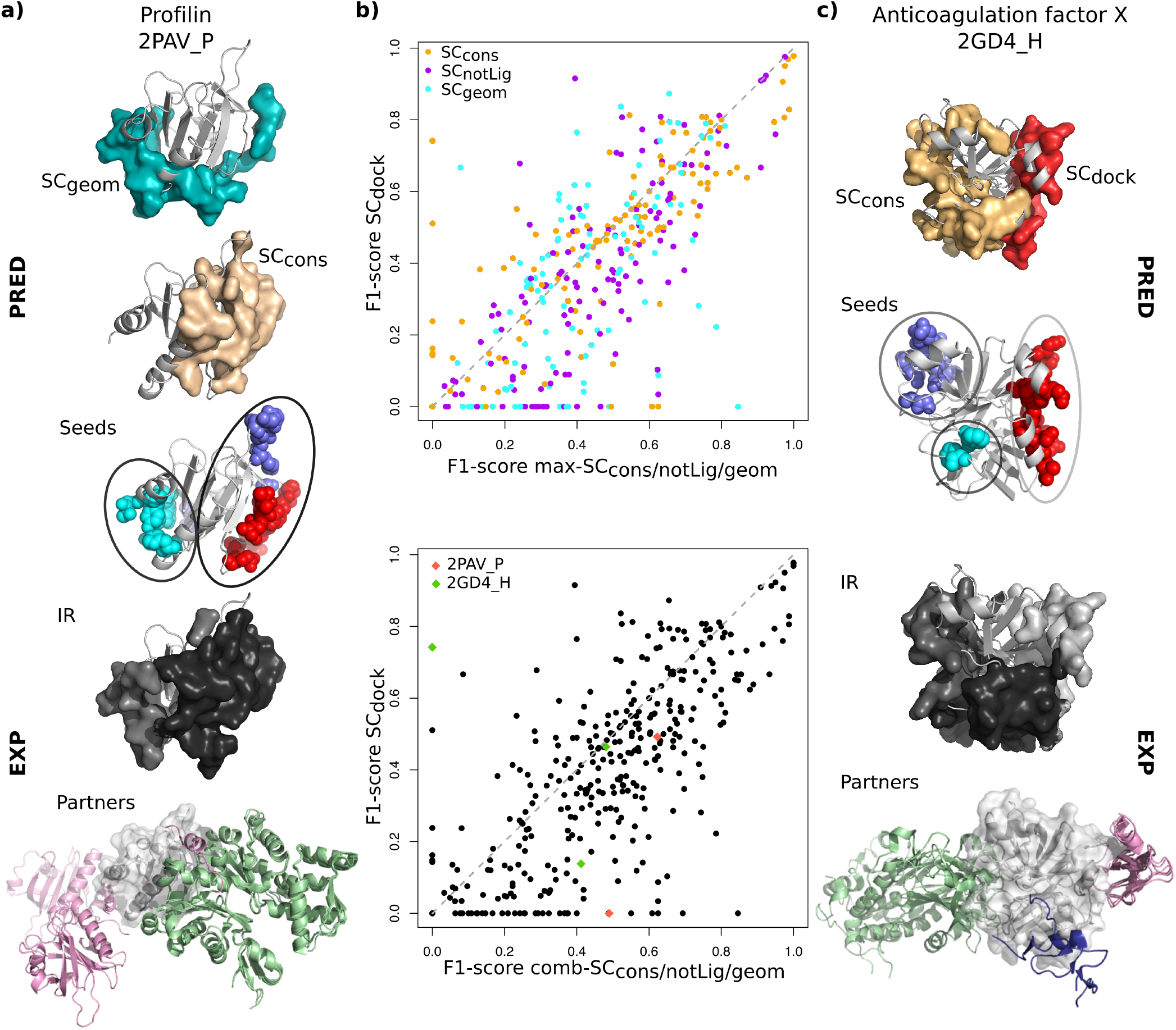
Examples and comparison of predictions. (a) Profilin (light grey cartoon) displayed with the patches predicted by SC_*cons*_ (in beige) and SC_*geom*_ (in cyan), the patches’ clustered seeds, two experimental IRs from PPI-262_ext_ (in grey tones) and the corresponding partners (colored cartoons); (b) Scatterplot of F1-scores computed for the best-matching patch or combination of patches, among SC_*cons*_, SC_*notLig*_, SC_*geom*_ (x-axis) and from SC_*dock*_ (y-axis) against experimental IRs from PPI-262_ext_. In cases where a combination of several patches is retained, the patches either come from a single SC (on top) or from several SC (at the bottom, x-axis). (c) Heavy chain of the anti-coagulation factor X (light grey cartoon) displayed with the patches predicted by SC_*cons*_ (beige) and SC_*dock*_ (red), the patches’ clustered seeds, the three experimental IRs from PPI-262_ext_ (in grey tones) and the corresponding partners (colored cartoons).

To better characterize the contribution of docking-based information, we compared the predictive performance of SC_*cons*_, SC_*notLig*_, SC_*geom*_, either considered individually or altogether, with that of SC_*dock*_ (Fig. 5b). We observed that the vast majority of IRs is better detected by the former than the latter (points below the diagonal, 68% on top and 72% at the bottom). Hence, evolutionary conservation, physico-chemical properties and local geometry are generally able to better capture protein interface signals than the coarse-grained empirical energy function used in the docking experiment. Nevertheless, there are a number of cases where docking-based data provide valuable information to improve predictions by unveiling interfaces that could not be detected otherwise. An example is given by the anticoagulation Factor X (Fig. 5c), where one of its three IRs (in white) is very well detected by SC_*dock*_ (in red, F1-score = 0.74) but completely missed by the other SCs.

### Predictions capture interface variability coming from molecular flexibility

Accurately accounting for molecular flexibility remains a challenge for protein interface and interaction prediction. We looked at how our predicted patches matched experimental interfaces undergoing variations from one structure to another. We focused on the 78 IRs from PPI-262_ext_ which are only slightly (<1.5 times) bigger than the corresponding IS(s) from `PPI-262`. Four examples are illustrated on Figure 6. The difference between the IR and the original IS(s) typically reflects the interface variability between different crystallographic structures of the same complex. For example, about 20 structures of the same hetero-4-mer involving Caspase-1 are available in the PDB and contribute to the definition of one IR (Fig. 6, top right). For the vast majority of these IRs (>85%), the precision reached by dynJET^2^ predictions is equal to or greater than that computed on the corresponding ISs (compare black/white and colored surfaces on Fig. 6). These results reveal that there exists a non-negligible variability inherent to protein interfaces and that dynJET^2^ predictions is generally able to capture it.

**Figure 6:**
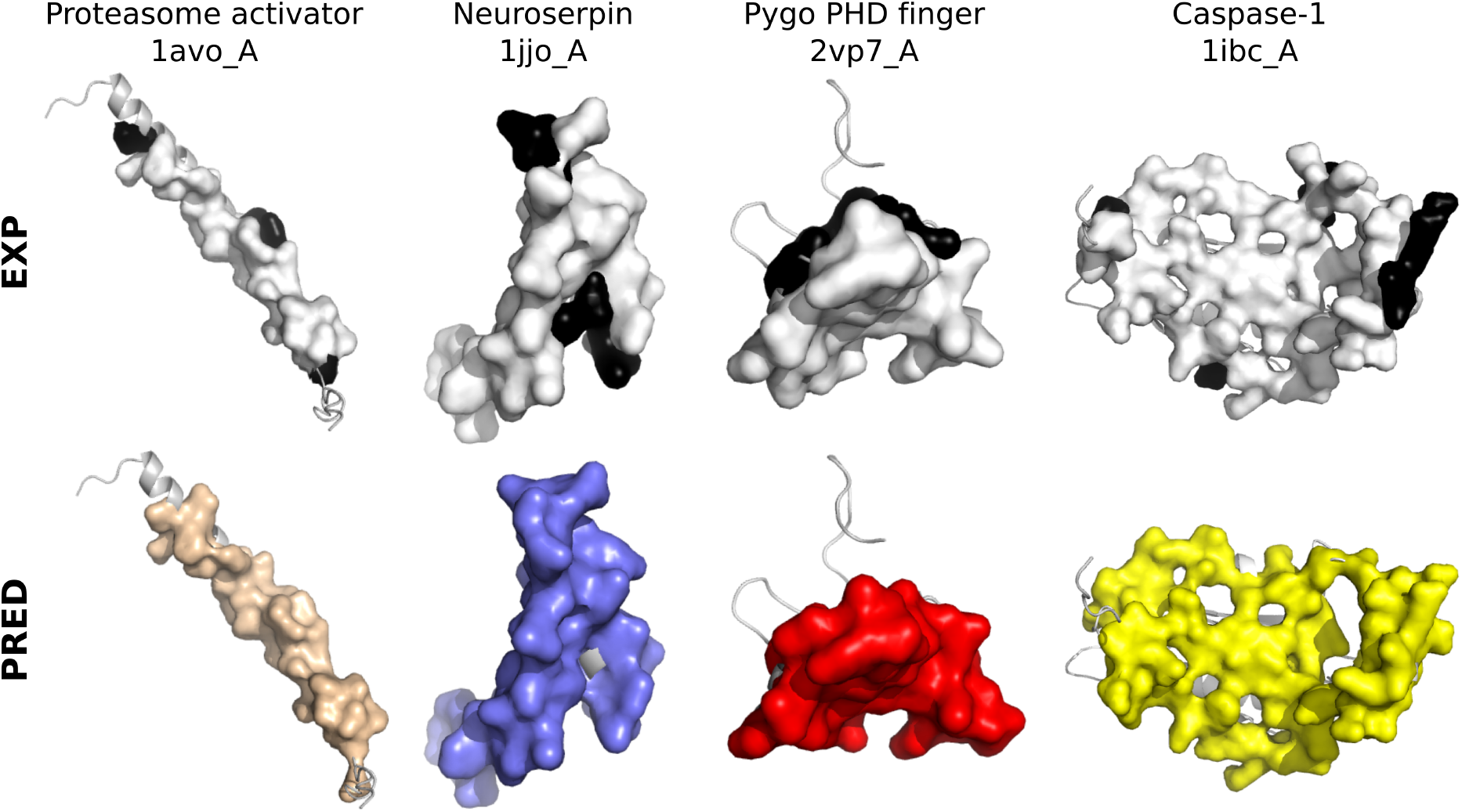
Examples of predictions whose precision is higher on the IR compared to the IS. The query protein structure from P-262 is displayed as a grey cartoon. The experimental and predicted interfaces are displayed as opaque surfaces: on top, the IS is colored in white and the additional residues belonging to the IR are in black; at the bottom, the SC_*cons*_, SC_*notLig*_ and SC_*dock*_ patches predicted for 1avo_A, 1jjo_A, 2vp7_A are in wheat, purple and red, respectively, and the best combination of patches predicted for 1ibc_A is in yellow. The precision increases from 79 to 91% for 1avo_A, from 76 to 92% for 1jjo_A, from 70 to 83% for 2vp7_A and from 75 to 84% for 1ibc_A.

We also assessed the robustness of our predictions with respect to conformational changes. For each IR from PPI-262_ext_, we calculated the conformational deviation of its backbone atoms between the query structure from `P-262`, on which our predictions were computed, and the structures of its homologs (see Materials and Methods). Almost all (95%) IRs display average conformational deviations lower than a 4A (S3 Fig). The extent of the deviation is not correlated to the quality of the predictions (Pearson correlation coefficient of 0.05 between RMSD and F1-score, S3 Fig). This indicates that dynJET^2^ predictions are robust to small to medium conformational changes.

### Predicted patches’ seeds describe the multiplicity of interactions

Almost all (94%) IRs from PPI-262_ext_ were detected, at least partially, by considering predictions issued by all four SCs (Fig. 4g). Some predicted patches display a good or very good match with a single IR. For example, the interface between profilin and human VASP (Fig. 5a, in black) is very well detected by SC_*cons*_ (Fig. 5a, in beige, *Sens* = 0.63, *PPV* = 0.61). Another example is given by the interface between the heavy and light chains of the anticoagulation factor X which is well detected by SC_*dock*_ (Fig. 5c), in red, Sens = 0.82, PPV = 0.68). Some other patches cover several IRs, as exemplified by SC_*geom*_ in Profilin (Fig. 5a, in cyan) and SC_*cons*_ in the factor X’s heavy chain (Fig. 5c, in beige). These cases are ambiguous if one considers a single SC. However, by crossing the information coming from different SCs, one may infer the existence of several IRs and thus resolve the ambiguity. For instance, the presence of a SC_*cons*_ patch partially overlapping with the SC_*geom*_ patch at the surface of Profilin (Fig. 5a) could be used as an indicator of the existence of 2 IRs and of the fact that the SC_*geom*_ patch extends over these 2 IRs.

To test whether this type of reasoning could be generalized, we systematically investigated how predicted patches were distributed over experimental IRs. For this, we explicitly considered the patches’ seeds, which are the first groups of residues being detected by dynJET^2^ clustering algorithm. We collected all seeds generated by SC_*notLig*_, SC_*geom*_ and SC_*dock*_ and clustered them based on 3D proximity (note that SC_*cons*_ seeds were not considered for this analysis, see Materials and Methods). The total number of resulting clustered seeds is 562, which corresponds to 2.14 seeds per protein on average. By comparison, the average number of IRs is 1.4. About one quarter of the seeds are completely inside an IR (100% precision) and almost than half of the seeds detect an IR with very high (≥80%) precision (Fig. 7a). In the examples of Profilin and factor X, the number of seeds is equal to the number of IRs and each seed points to a different IR (Fig. 5a,c).

**Figure 7:**
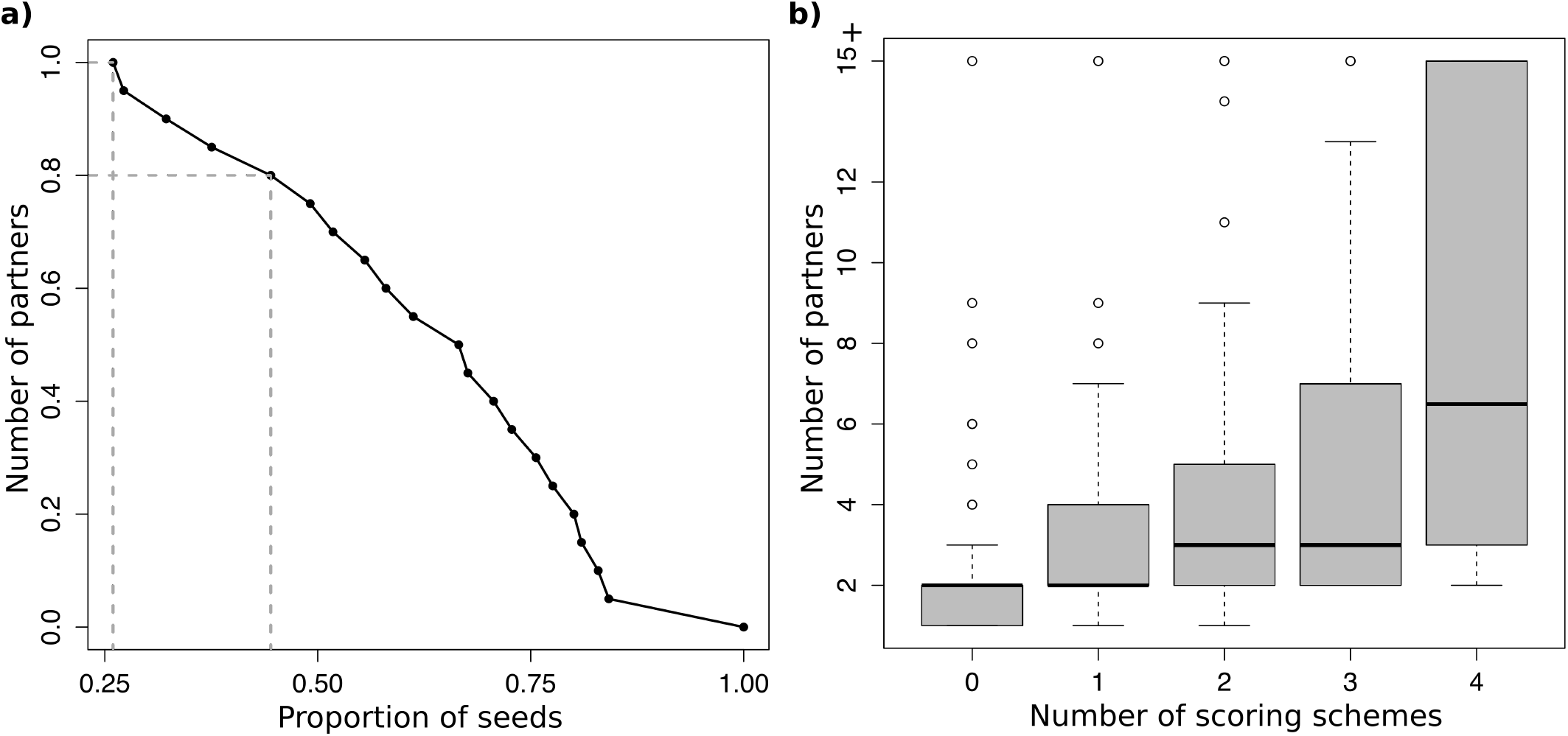
Ability of the patches’ seeds to detect IRs and estimate the number of partners. (a) Cumulative distribution of patches’ seeds precision in detecting IRs. Each x-value corresponds to the proportion of seeds with precision higher than the y-value. Dotted segments emphasize the points with *y* = 1 and *y* = 0.8. (b) Number of partners for each IR versus number of scoring schemes predicting a seed in the IR.

We also investigated whether seeds could be used to infer the number of partners a protein has (Fig. 7b). For this, we looked at the properties of the seeds lying completely or almost completely (PPV≥80%) within an IR. We observed that the number of partners binding to an IR increases with the number of scoring schemes predicting one or more seeds within the IR (Fig. 7b, Pearson correlation of 0.52). This means that IRs displaying a multiplicity of signals relevant to protein interactions tend to attract more partners. Hence, the accumulation of seeds with different properties in a protein region can be used as an indicator that this region will likely interact with many partners.

### Comparison with another state-of-the art interface predictor

We compared dynJET^2^ predictions to those of Multi-VORFFIP [50], a state-of-the-art machine learning method, integrating a broad set of residue descriptors including solvent accessibility, energy terms, sequence conservation, crystallographic B-factors and Voronoi Diagrams-derived contact density, in a two-steps random forest ensemble classifier. The distribution of F1-scores obtained for the union of the residues predicted by Multi-VORFFIP against a subset of PPI-262_ext_(see Materials and Methods) is similar to that obtained for the union of dynJET^2^predictions (Fig. 4a, compare the second and third boxes). However, the performance of Multi-VORFFIP in detecting individual IR is lower than that of dynJET^2^ (Fig. 4bcd, compare darkblue and green boxplots). The average F1-score is lower, the number of very good predictions is significantly lower and the number of missed IRs is much larger (52 versus 21 for dynJET^2^). In particular, our predictions are more precise than those of Multi-VORFFIP (Fig. 4d).

## Discussion

Protein surfaces are used in multiple ways in cellular partners association. A comprehensive and accurate description of protein surfaces should account for multiple partners, molecular flexibility (from slight rearrangements to conformational changes), disorder and post-translational modifications. In this work, we have analyzed a pool of proteins with different functions to address the two first aspects.

In line with a previous study [12], we found that the protein surface involved in functional interactions is probably much bigger than anticipated. An accurate estimation of this surface is mandatory for the correct assessment of protein interface prediction methods. However, such an estimation is still beyond reach as we do not know the exact number of cellular partners a protein has and how these partners use its surface in solution. To move forward, we have introduced the notion of interacting region, which results from combining several overlapping interacting sites detected in experimental complex structures. By taking into account all known homologs of our query proteins and their crystallographic complexes, we could synthesize over 23 000 ISs into a relatively small number of IRs (1.4 per protein chain). This procedure permitted to shed light on the variability of binding modes and on the multiplicity of protein surface usage. In the evaluation of dynJET^2^ predictions, we could appreciate that a large amount of predicted patches better matched IRs, compared to ISs. This result is expected from a good protein interface prediction algorithm, as the notion of IR seems more biologically pertinent than that of IS in many cases, especially when the IR synthesizes the variability inherent to structure ensembles of the same complex.

Our predictions were generated based on three sequence- and structure-based properties of monomeric protein surfaces and also on residue propensities inferred from docking calculations. The latter reflect how energetically favorable the interatomic interactions established between a protein and many other proteins are. The energy was evaluated by a combination of Lennard-Jones and Coulomb potentials. We have systematically assessed the contribution of this physical description of protein interactions to the prediction of interfaces. We have shown that in most cases, its predictive power is lower than that of the sequence and structure-based descriptors. Nevertheless, it brings complementary information which helps improving the accuracy of the predictions and, in some cases, it even unveils binding sites that could not be detected otherwise.

We have highlighted several cases where almost the entire protein surface is involved in functional interactions. This finding challenges the role of specificity in the evaluation of protein interface prediction methods and rather put the emphasis on precision. We have demonstrated the usefulness of the prediction patches’ three-layer structure by showing that the patches’ seeds enabled precisely locating and discriminating IRs at the protein surface. We have further shown that the seeds could help determine whether a protein has a few or many partners. Future work will aim at getting a more accurate estimation of the number of partners. Moreover, with the help of future PPI data, it seems achievable to associate functions to the partners binding on different surface areas, described by different seeds on a region.

Although dynJET^2^ predictions match reasonably well experimentally identified interacting regions, the match is not perfect. Given the degree of complexity we have highlighted in the usage of a protein surface, it seems legitimate to ask whether perfect match with experimental interfaces is an attainable goal for protein interface predictions algorithms that, like dynJET^2^, do not use any knowledge about the query protein’s partners. An accurate estimation of the maximum level of agreement one could expect would be most valuable. Besides, even without perfect match, dynJET^2^ predictions can be fully exploited to guide experiments. For example, the above-mentioned ability of patches’ seeds to precisely locate IRs has implications for the control and modulation of existing protein-protein interactions. Mutating seed residues should impact the binding of the associated partners. Another way to go would be to design interactors that bind to the predicted patches. Indeed, dynJET^2^ algorithm provides a mean to delineate regions at the protein surface with sizes similar to those of experimental interacting sites or regions and complying with a small number of properties known to be relevant to protein-protein association. Thus, in addition to detecting regions being actually used to interact, it also reveals the potentiality of other regions to interact.

## Materials and Methods

### Datasets

#### Proteins: P-262

We defined a dataset of 262 protein chains and associated PDB structures featuring both single and multiple partners interactions. This dataset was extracted from a larger set of 2246 protein chains defined in the scope of the HCMD2 project (see http://www.ihes.fr/~carbone/HCMDproject.htm), for which we performed Complete Cross-Docking. We considered the subset of PDB structures comprising a protein complex previously reported in [45], from which we excluded: (a) only C-α structures, (b) chains for which docking results were missing, (c) chains forming coiled-coils complexes, (d) deprecated PDB codes, (e) chains for which no biologically relevant interface (see Methods for definition) could be found in the whole PDB (considering 90% sequence identity, see Methods). The remaining 262 protein chains comprise on average 200.5 ± 131.2 residues (S1 Table). This indicates a large variation of protein size inside the dataset (21 residues for the smallest protein versus 789 residues for the largest one).

#### Protein interfaces: PPI-262 and PPI-262_ext_

We defined two datasets of experimental protein-protein interfaces, namely `PPI-262` and PPI-262_ext_ (S1 Fig). Both datasets comprise only interfaces buried within ‘‘biological units” or ‘‘biological assemblies”, as annotated by the authors of the PDB structure or by PISA software [51]). This ensures that the interfaces we consider carry a biological meaning. `PPI-262` comprises 329 ISs (see definition below, in Methods) extracted from the PDB files associated to `P-262` and PPI-262_ext_ comprises 370 IRs (see definition below, in Methods) defined from PDB files of close homologs of the proteins from `P-262`.

To construct PPI-262_ext_(see S1 Fig), we first searched for homologs of the 262 proteins from `P-262`in the PDB. We downloaded the pre-computed set of PDB structures clustered at 90% sequence identity from ftp://resources.rcsb.org/sequence/clusters/. This set was determined using `BLASTClust` with the arguments `-p T -b T -S 90`. We then filtered out structures with a resolution poorer than 5 Å resolution. 23 642 functional ISs were detected on these structures and were then mapped onto the query proteins from `P-262` by performing global pairwise sequence alignment (using the `blosum62` matrix, with the Biopython package [52]). ISs were then merged into IRs.

#### Complete Cross-Docking

Given an ensemble of proteins, Complete Cross-Docking (CC-D) consists in docking each protein against all others in the dataset, including itself. CC-D was performed on `P-262` using the MAXDo (Molecular Association via Cross Docking) rigid-body coarse-grained docking program [42]. Statistics were computed from the generated conformations (docking poses) to determine the propensity of each residue from each protein to be found in a docked interface. We define the interface propensity (IP) of residue i, belonging to protein *P*, as [42, 11]:

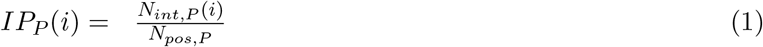

where *N_pos_,p* is the total number of docking poses considered for protein *P* and *N_int,p_*(*i*) is the number of docking poses where residue *i* lies in the interface. In order to limit the number of docked interfaces to reconstruct, which is the most time-consuming part of the analysis, we considered only the lowest energy docking poses (less than 2.7 kcal from the best-scored pose, as described in [11]). This led to approximately 50 000 docking poses for each protein pair. Thus, for a given protein *P*, we considered about 50 000 × 262 = 13 100 000 docking poses.

Below, we shall use *IP_P_*(*i*) values to formally define a normalised interaction propensity score, called NIP, that dynJET^2^ uses in order to predict interface sites.

#### Residue-based properties

Four measures, T_*JET*_, PC, CV and NIP, are used to evaluate single residues in a protein and to define scores for the prediction of protein interfaces.

T_*JET*_ reflects the evolutionary conservation level of a residue, and is computed from phylogenetic trees constructed by using sequences, homologous to a query sequence and sampled by a Gibbs-like approach [23]. The Gibbs-like approach extracts *N* representative subsets of *N* sequences [23] in a way that, within each subset, the proportions of sequences sharing [20 – 39]%, [40 – 59]%, [60 – 79]%, and [80 – 98]% sequence identity with the query sequence are similar (ideally, about one quarter for each group of identity). Sequences in a subset are then aligned using CLUSTALW2 [53] and a distance tree is constructed from the alignment based on the Neighbor Joining algorithm [54]. From each tree *T, a tree trace level* is computed for each position in the query sequence: it corresponds to the level n in the tree *T* where the amino acid at this position appeared and remained conserved thereafter (see [23] for a more precise definition). Let us recall that this definition of evolutionary trace is notably different from the measure defined in [14, 55] to rank protein residues.

Then, tree trace levels are averaged over the *N* trees to get statistically significant values, which we denote *relative trace significances*, or *T_JET_*, and which are calculated as follows [23]:

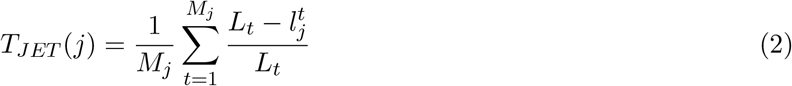

where 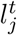 is the tree trace level of residue *r_j_* in tree *t, L_t_* is the maximum level of *t* and *M_j_* is the number of trees where a non-zero tree trace level was computed for *r_j_*. *T_JET_* values vary in the interval [0,1] and represent averages, over all trees of residues, of evolutionary conservation levels.

PC indicates the physico-chemical propensity specific to amino acids located at a protein interface. The original values, taken from [56], range from 0 to 2.21 and are scaled here between 0 and 1 for the calculation of residue scores.

CV is the circular variance, a measure of the vectorial distribution of a set of neighbouring points around a fixed point in 3D space [57]. For a given residue, CV reflects the density of the protein around it: residues buried within the protein will display high CV values, while exposed or protruding residues will display low CV values. Compared to solvent accessibility, CV changes more smoothly from the surface to the interior of the protein [58], and is thus less sensitive to small conformational changes. CV can be applied equally well to atomic or coarse-grain representations [57]. The CV value of an atom *i* is computed as:

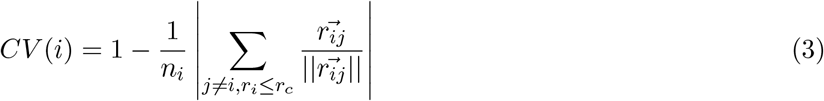

where *n_i_* is the number of atoms distant by less than *r_c_*Å from atom *i*. The CV value of a residue *j* is then computed as the average of the atomic CVs, over all atoms of *j*. A low CV value indicates that a residue is located in a protruding region of the protein surface. CV values are scaled between 0 (most protruding residues) and 1 (least protruding residues) for the calculation of residue scores.

NIP is the normalised form of the Interface Propensity score *IP*, defined in Eq. (1), that reflects the propensity of a residue to be found at the interface. In order to compare *IP* scores among proteins, we normalise it, as done in [11]: a positive NIP value indicates that the residue *i* is favoured to occur at potential binding sites, and a negative NIP value indicates that it is disfavoured. NIP is defined as:

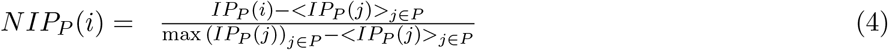

where < *IP_P_*(*j*) >_*j*∈*P*_ and max(*IP_P_*(*j*))_*j*∈*P*_ are the average *IP* and the maximum *IP*, respectively, computed over all the residues *j* in *P*. The NIP value represents how often a residue is docked on the retained conformations (that is, those conformations that have less than 2.7*kcal*/*mol* energy difference from the best one, as explained above).

These four residue-based properties were previously shown to be useful for the prediction of protein interfaces [33, 44, 43, 24, 11, 59, 23]. T_JET_, PC, CV are computed using dynJET^2^, a modified version of JET^2^ [24] that handles NIP values, as described below. For each measure, values are scaled between 0 and 1.

#### Surfaces and interfaces

A residue is considered to be at the surface of the protein if it displays at least 5% of relative accessible surface area (rasa), as computed by Naccess [60]. Experimental and predicted interfaces are exclusively comprised of surface residues.

*Experimental interfaces* are detected on known PDB complex structures, considering only biological assemblies. We define two types of interfaces, namely *interacting sites* (ISs) and *interacting regions* (IRs). ISs are detected in single PDB structures by using the INTBuilder software [61] (www.lcqb.upmc.fr/INTBuilder/) with a distance threshold of 5Å. They may represent binary interactions or multiple partners interactions. As an example, let A, B and C be 3 proteins involved in a complex where B and C both bind to A. If B is less than 5Å away (minimal distance) from C, then we consider proteins B and C to be in contact and describing a multi-partner interaction with A. Otherwise, we consider the 2 binary interactions A-B and A-C separately. ISs smaller than 6 residues were discarded for the analysis. IRs are defined by merging several ISs. Two ISs, namely *IS*_1_ and *IS*_2_, are merged into an IR if their maximum overlap with respect to their respective sizes (max{overlap(*IS*_1_, *IS*_2_), overlap(IS_2_, IS_1_)}) is greater than an arbitrarily chosen threshold of 60%. In practice, this threshold gave us the most realistic IRs given the experimental information. Additionally, any IS of at most 5 residues will be merged with another IS if the two ISs have at least one residue in common. The merging procedure is iterated over all ISs for a given protein.

*Predicted interfaces* are identified by the dynJET^2^ software (www.lcqb.upmc.fr/dynJET2/). Given the sequence and the structure of a query protein, dynJET^2^ predicts the location of potential protein binding sites on the protein surface [24]. dynJET^2^ implements a clustering algorithm and scoring strategies specifically aimed at detecting the different layers of a protein interface, as defined in [48]. The algorithm [24] first identifies a small cluster of highly scored residues, called the seed. Seeds closer than 5 Å are merged. Then, the detected seeds are progressively extended, and the resulting residue clusters are merged if they are in contact (< 5 Å away). Finally, an outer layer is added to form what we call a *predicted patch*. We used the iterative mode of dynJET^2^ (i-dynJET^2^) and considered a residue to be predicted as interacting if it was detected at least twice over 10 iterations (as done in [24]).

Four scoring schemes or strategies are implemented in dynJET^2^ (compared to three in JET^2^ [24]):

**SC**_*cons*_ targets very conserved residues (identified by the T_*JET*_ score) to form a seed which is then extended using both T_*JET*_ and PC scorings. An outer layer is added considering both PC and CV scorings. SC_*cons*_ is intended to detect diverse protein binding sites. This step, essentially unchanged compared to the original JET version, was extensively described in [23].
**SC**_*notLig*_ detects both seed and extension layers using a combination of T_JET_ and CV scorings. It aims at detecting highly conserved residues that are not buried too deeply beneath the surface of the protein. The outer layer is defined based on PC and CV, as in SC_*cons*_. SC_*notLig*_ specifically distinguishes protein interfaces from small ligand binding sites.
**SC**_*geom*_ disregards evolutionary information and solely employs PC and CV for detecting all three layers of the interface. SC_*geom*_ yields consistent predictions for interfaces displaying very low conservation signal, *e.g*. antigen binding sites.
**SC**_*dock*_ applies the NIP score of the residues to all three layers (core, extension and outer layer). The usage of NIP is motivated by the observation that proteins tend to dock to their cognate partners and also to non-interactors via the same region at their surface [42, 59, 11, 43]. This scoring scheme is new in dynJET^2^.

To evaluate the performance of our predictions, we mainly relied on the F1-score, which is the harmonic mean of precision and recall. Predicted patches were compared to ISs from `PPI-262` and IRs from PPI-262_ext_. The union of all predicted interface residues was also compared to the union of all experimental interface residues, for each protein.

#### Seeds clustering

Seeds generated by dynJET^2^’s different scoring schemes were collected. The SC_*cons*_ seeds were discarded because almost half of their residues were shared with the SC_*notLig*_ seeds (S2 Figa), they were bigger than the other seeds (S2 Figb) and we observed that they often extended over several other seeds. We considered that these characteristics would make SC_*cons*_ seeds perform badly in locating different IRs. The atoms belonging to the seeds collected from SC_*notLig*_, SC_*geom*_ and SC_*dock*_ were then classified by applying hierarchical clustering using the average linkage method. A threshold distance of 23Å was used to define the clusters. This value yielded the best match between the number of clustered seeds and the number of IRs.

#### Conformational variability of IRs

For each IR, the RMSD of its backbone atoms (or, if not possible, its C-α atoms) was computed between the query structure from `P-262` and each of the homologous structures on which the IR was detected. For each homologous structure, only the subset of residues detected on this structure were considered to compute the RMSD. RMSD values were then averaged over the homologous structures (including the query structure if the IR was also detected on it). This gives us a single RMSD value for each IR.

#### Number of partners

To count how many different partners a protein has, we considered all known homologs of the protein in the PDB and their partners. We clustered the partners depending on their sequence homology: two partners were classified in different clusters if they shared less than 90% sequence identity. This threshold in agreement with the criteria we applied to protein chains and their homologs. The number of clusters provides an estimation of the number of partners for the protein.

#### Comparison with Multi-VORFFIP

Multi-VORFFIP [50] (accessed at ^1^) was applied on 252 protein chains from `P-262`. The 10 proteins that were discarded for the analysis either belonged to the training set of Multi-VORFFIP or produced an error when running the tool. We considered residues displaying a probability above 0.5 as predicted to interact. Residues separated by less than 5A were clustered to form predicted patches.

## Acknowledgements

The MAPPING project (ANR-11-BINF-0003, Excellence Programme “ Investissement d’Avenir”); funds from the Institut Universitaire de France; the access to the HPC resources of the Institute for Scientific Computing and Simulation (Equip@Meso project - ANR-10-EQPX-29-01, Excellence Program “ Investissement d’Avenir”); the World Community Grid (`WCG`, www.worldcommunitygrid.org) and WCG volunteers that allowed us to perform cross-docking experiments with MAXDo on the HCMD2 dataset.

## Supporting Information Captions

**S1 Table**

**List of proteins in the** `P-262` **dataset**.

**S1 Fig**

**Schematic representation of the protocol applied to collect interacting sites and regions**.

**S2 Fig**

**Characteristic features of the seeds detected by dynJET^2^**. (a) Overlap between seeds generated by the different dynJET^2^ scoring schemes, averaged over all proteins from `P-262`. The overlap is computed as: 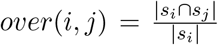, where *s_i_* is the ensemble of seed residues predicted by the *i^th^*scoring scheme. (b) Distributions of the sizes (in residues) of the seeds predicted by dynJET^2^ different scoring schemes.

**S3 Fig**

**Representation of the RMSD values compared to the F1-scores**. Each point represents the values obtained for one IR. The RMSD values are computed on the backbone atoms between the query structure from `P-262` and each of the homologous structures on which the IR was detected. The F1-score values correspond to the best combination of dynJET^2^ predictions.

## Footnotes

1 www.bioinsilico.org/cgi-bin/SUPER_VORFFI/htmlVORFFI/home

